# Mapping of individual somatosensory representations – comparison of fMRI and TMS

**DOI:** 10.1101/2025.06.12.659279

**Authors:** Juha Gogulski, Mikko Nyrhinen, Rasmus Zetter, Maksym Tokariev, Hai Lin, Antti Pertovaara, Synnöve Carlson

**Affiliations:** Department of Physiology, Faculty of Medicine, University of Helsinki, Helsinki 00014, Finland; Department of Neuroscience and Biomedical Engineering, Aalto University School of Science, Aalto 00076, Finland; Aalto TMS, Aalto NeuroImaging, Aalto University School of Science, Aalto 00076, Finland; Department of Neurosurgery, Shenzhen Second People’s Hospital, the First Affiliated Hospital of Shenzhen University, Shenzhen, China

## Abstract

Invasive brain mapping, functional magnetic resonance imaging (fMRI) and navigated transcranial magnetic stimulation (nTMS) results indicate that somatosensory representations vary between individuals. However, it is unknown how well somatosensory representations determined using nTMS and fMRI correspond. Here, we used single-pulse nTMS and fMRI in 17 right-handed subjects to determine the S1 representation of the tip of the right index finger stimulated mechanically with a Braille device. In an nTMS mapping experiment, the S1 site at which tactile sensation was blocked by nTMS, was considered the S1 representation site (S1_HS_) of the fingertip. The location of S1_HS_ varied up to 36 mm between subjects. In the fMRI experiment, passive and oddball tactile tasks were employed. The location of the somatosensory peak fMRI activation varied up to 39 mm between subjects in the passive condition and up to 84 mm in the oddball condition. Within subjects, the mean distance between the S1_HS_ and the peak fMRI activation was 10 + 2 mm (S.E.M.) in the passive condition and 14 + 3 mm in the oddball task. Taken together, our findings underscore the importance of multimodal approaches to brain mapping in future studies.

## 1. Introduction

Intact somatosensory pathways are fundamental for both sensory perception and motor control (Okuda et al. 1995; Tang et al. 2023). The primary somatosensory cortex (S1) represents a critical node in processing tactile information, yet our understanding of individual variations in these representations remains limited. Accurate mapping of somatosensory cortical areas has significant implications for both basic neuroscience research and clinical applications, particularly in presurgical planning. While various mapping techniques exist, each offers distinct advantages and limitations that must be carefully considered when interpreting functional localization results.

Somatotopy of the S1 has been non-invasively studied by various investigators using functional magnetic resonance imaging (fMRI) (van Westen et al. 2004; Martuzzi et al. 2014; Puckett et al. 2017). Single-pulse transcranial magnetic stimulation (TMS) provides another non-invasive method for temporally and spatially accurate mapping of somatotopic organization of the S1, particularly when TMS is connected to a navigation system (nTMS). When compared with correlative data obtained with fMRI, nTMS has the additional benefit that it can induce blocking of the studied somatic sensation in a highly reproducible manner (Hannula et al. 2005), thereby giving information about the causal relationship of the sensation evoked by the cutaneous test site stimulation with the S1 target of nTMS. So far, nTMS has been rarely used in S1 mapping (Hannula et al. 2005; Gogulski et al. 2015; Gogulski et al. 2017; Tang et al. 2023). It is not yet known how closely cortical somatosensory representations determined with nTMS correspond with those determined with fMRI. Moreover, earlier fMRI (van Westen et al. 2004; Martuzzi et al. 2014) and nTMS (Gogulski et al. 2017) studies as well as invasive mapping results (Penfield and Boldrey 1937) indicate that there is marked variability between individuals in the locations of somatic presentations. The correspondence between inter-individual variability in the S1 representations obtained with fMRI and nTMS has not yet been quantitatively studied.

Here we used single-pulse nTMS and fMRI to determine inter-individual variability in the location of the somatosensory representation of the tip of the right index finger that was stimulated mechanically with a Braille device. The S1 representation site (S1_hotspot = HS_) was defined as the S1 site at which nTMS blocked the sensation elicited by the mechanical stimulus. We used passive tactile stimulation and a tactile oddball task to determine somatosensory representation area with fMRI. We hypothesized that 1) the location of the S1_HS_ would show significant inter-individual variability; 2) stimulation of S1_HS_ would impair the spatial discrimination ability; 3) within individuals, location of the S1_HS_ would be consistent with fMRI activation evoked by tactile stimulation; 4) variability of the S1_HS_ locations across subjects would be similar compared to variability of fMRI peak activation locations. Our results showed that 1) the location of the S1_HS_ varied considerably across individuals; 2) stimulation of S1_HS_ did not affect the spatial discrimination ability; 3) the fMRI activation evoked by tactile stimulation and the nTMS-defined S1_HS_ were not always overlapping; 4) across subjects, the variability of fMRI peak activation locations was larger than the variability of S1_HS_ locations. Discrepancies between fMRI and nTMS results suggest that fMRI may reflect a broader activation area including both direct and indirect somatosensory processing networks, while nTMS is a more focused method pinpointing the functional hotspot critical to tactile detection.

## 2. Methods

### 2.1. Subjects

Seventeen healthy, right-handed volunteers (mean age 29 years, age range 24-37 years, 8 females) with no history of neurological or psychiatric disorders participated in the study. All subjects participated in the nTMS blocking experiment, three of which were excluded due to failure to find an S1 site that would block tactile perception. Eleven subjects (mean age 29 years, age range 26-36 years, 5 females) participated in the spatial discrimination experiment. Ten of the 17 subjects (mean age 30 years, age range 26-36 years, 4 females) participated in the fMRI experiment. Informed consent was obtained from all participants. The study was conducted according to the World Medical Association’s Declaration of Helsinki and approved by the Aalto University Ethics Committee.

### 2.2. Tactile stimulator

Tactile stimuli were applied to the fingertip of the right index finger using a piezoelectric stimulator unit (Metec AG, Stuttgart, Germany) originally designed for Braille reading, which was driven by a custom-made control system. The stimulator included 8 stimulus probes in a 2×4 grid configuration. Each stimulus probe was a rounded plastic pin (diameter 1.25 mm) that can be raised to a height of 0.7 mm with a rise time of 24 ms, generating a force of approximately 0.17 N. In the nTMS blocking experiment, stimulus amplitude was controlled by varying the rise time, whereas in the fMRI experiment and in the spatial discrimination experiment, a stimulus duration of 50 ms was used. 50 ms stimulus was chosen to be sure that all stimuli are being perceived. Optimal stimulus parameters were explored in a preliminary experiment (Supplementary Fig. 1).

### 2.3. Navigated TMS

Anatomical, T1-weighted MR-images were obtained with a 3T MRI scanner before the nTMS experiments. TMS sessions were conducted at the Aalto TMS laboratory (Aalto NeuroImaging, Aalto University School of Science) using a Magstim 200^2^ stimulator unit with a Magstim 70 mm figure-of-eight coil (P/N 9925-00; Magstim Co., Carmarthenshire, UK), Visor2 neuronavigation system (ANT Neuro, Enschede, Netherlands), and monophasic single pulses. Presentation software (Neurobehavioral Systems, Albany, CA, USA) was used to control the delivery of the tactile and nTMS stimuli and to log the subjects’ responses. The control TMS site was defined as a midline site located 3 cm above the inion, corresponding to the Oz electrode position of the 10–20 electroencephalography (EEG) electrode positioning system (Schutter and van Honk 2006).

### 2.4. Blocking of tactile perception with navigated TMS

Figure 1a illustrates the workflow of the nTMS experiment. First, the resting motor threshold (MT) was determined for each participant. To determine the MT, we first located the area on the left primary motor cortex (M1) corresponding to the right abductor pollicis brevis (APB) muscle. This was accomplished by directing the electric field anteriorly and perpendicularly relative to the precentral sulcus. Motor evoked potentials from the APB were captured using the NeurOne electromyography system (Mega Electronics Ltd., Kuopio, Finland), with Ag-AgCl skin electrodes (Spes Medica Srl., Genova, Italy). The M1 site that elicited the strongest motor responses was identified as the M1 hotspot (M1_HS_) and was used to determine the MT. The MT was defined as the lowest TMS intensity at which ≥ 5 / 10 stimuli resulted in a peak-to-peak EMG response of ≥ 50 μV.

**Figure 1.**
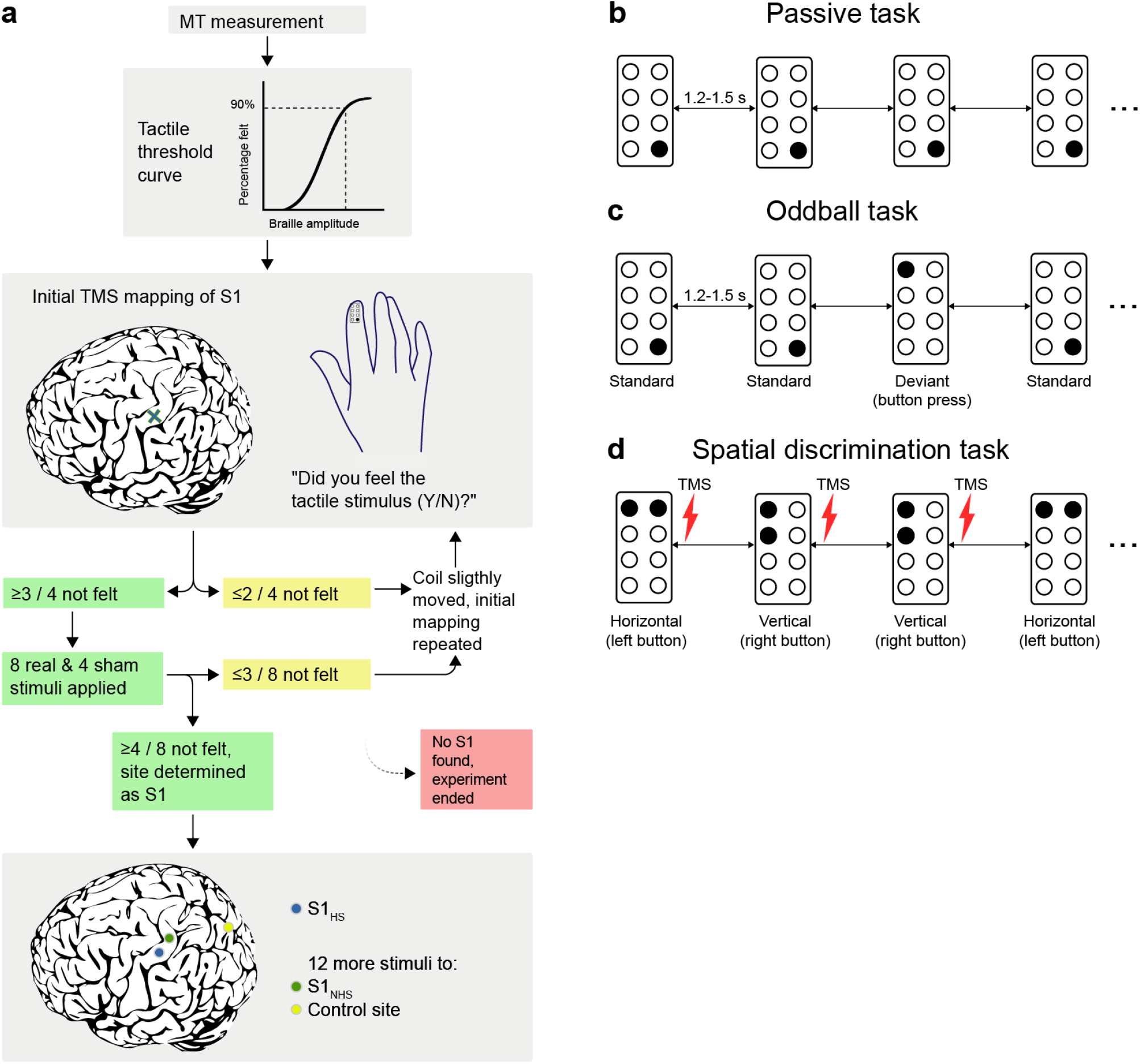
Experimental design of nTMS and fMRI mapping experiments. **a** Flow-chart of the nTMS experiment. MT = Motor threshold, S1_HS_ = S1 hotspot, S1_NHS_ = S1 non-hotspot. **b and c** fMRI localization of the right index finger representation. **b** In the passive tactile fMRI task, there was no button press required. **c** In the oddball task, the subject was instructed to press the button when the deviant stimulus was presented. **d** Spatial discrimination task (nTMS). The subject was instructed to discriminate between horizontal and vertical stimuli.

After MT assessment, an individual tactile threshold curve was assessed (Supplementary Fig. 1). Tactile stimulus amplitude which was felt in 90% of the trials was used for the nTMS experiment. Subjects were instructed to visually fixate on a cross that was displayed on a screen. To determine the S1 site that blocks perception of the tactile stimulus, we started to explore the S1 by turning the TMS coil 180° from the M1_HS_. A block of four trials was applied to the S1 location posterior to the M1_HS_ (initial mapping). Each trial consisted of a tactile stimulus and an nTMS pulse. TMS was applied at 120% of the resting MT except for three subjects, for whom the intensity had to be lowered due to uncomfortable scalp sensations (110% for one subject and 100% for two subjects). The average TMS intensity was 48% of the maximal stimulator output. Based on an earlier study (Hannula et al. 2005), the nTMS pulse was applied 20 ms after the onset of the tactile stimulus. If the subject felt ≥ 2 of the 4 stimuli, the coil was moved approximately 5 mm to another location on the postcentral gyrus. This was repeated until we found the S1 site where nTMS blocked the sensation of ≥ 3 of the 4 stimuli. For this site, a longer block of 12 trials was applied. Of these 12 trials, 8 included tactile stimuli and 4 were sham trials (only nTMS, no tactile stimulus). Sham and real tactile stimulus trials were presented in random order and nTMS was applied in all trials. The cortical site where nTMS blocked the perception of ≥ 50 % of the real tactile stimuli was defined as the S1 hotspot (S1_HS_). The mapping was continued until such a site was found. If after several attempts the S1_HS_ was not found, the experiment was ended.

We also examined the spatial extent of the somatotopic blocking effect of nTMS. For a subset of 11 subjects, the coil was moved along the postcentral gyrus to a distance of approximately 1 cm from the S1_HS_. This location was defined as the S1 non-hotspot (S1_NHS_).

The subjects responded by pressing the left computer mouse button with their left middle finger if they felt the stimulus, and the right mouse button with the left index finger if they did not perceive the stimulus. Subjects were instructed to press the button as quickly and as accurately as possible. After pressing the button, the subjects gave a verbal confidence rating for each response on a 3-level scale: a guess (= 1, low level of confidence), unsure (= 2, medium level of confidence), or sure (= 3, high level of confidence). The inter-trial interval, measured from the response, was 3000-3500 ms (jittered).

### 2.5. Influence of nTMS on spatial discrimination ability

Earlier studies showed that S1_HS_ is involved not only in the detection but also temporal discrimination of tactile stimuli (Hannula et al. 2005; Hannula et al. 2008), while the role of S1_HS_ in discriminating the spatial stimulus properties is not yet known. Thus, in an additional TMS experiment (N = 11), we explored whether nTMS of the S1_HS_ influences subjects’ tactile spatial discrimination ability (Fig 1d). The subjects were delivered Braille stimuli to the right index finger with two pins positioned either vertically or horizontally. By pressing the mouse buttons with the left hand, the subjects responded whether they felt a horizontal or a vertical tactile stimulus. TMS pulses were delivered 20 ms after the onset of tactile stimuli. In total, four blocks of 30 stimuli were presented (two blocks with TMS of S1_HS_ and two to the cortical control site). The order of blocks was counterbalanced between subjects.

### 2.6. MRI data acquisition

Functional and structural MRI data were acquired using a 3T MAGNETOM Skyra whole-body scanner (Siemens Healthcare, Erlangen, Germany) and a 30-channel receiving head coil. The subjects wore ear plugs and tightly packed foam covers over their ears for additional hearing protection and to minimize head movement. A T1-weighted 3D-MPRAGE structural image was obtained using the following parameters: TR 2530 ms, TE 3.3 ms, TI 1100 ms, flip angle 7°, FOV 256 × 256 mm, 1 mm isotropic voxels and 176 contiguous slices. Functional images were acquired using a gradient-echo echo-planar sequence with the following parameters: TR 2500 ms, TE 30 ms, flip angle 75°, FOV 220 × 220 mm, 3.4 mm isotropic voxels and 45 contiguous slices.

### 2.7. fMRI mapping procedure

The fMRI experiment was designed to localize the cortical somatosensory representation area of the tip of the right index finger at individual and at group level, thus allowing a comparison with the S1_HS_ sites obtained in the nTMS blocking experiment. The fMRI session was separated into the ‘passive’ and ‘oddball’ imaging runs, with short breaks in between runs.

First, the subjects performed a passive condition (hereafter called the passive task), in which they were told that they will receive tactile stimuli to the tip of their index finger (Fig. 1b). During the first five passive blocks, stimuli were delivered to the left index finger (30 stimuli per block; interstimulus interval of 1200-1500 ms; randomized, 10 ms step size). Then, similar tactile stimulation blocks were delivered to the right index finger. The blocks were separated by four rest blocks with a duration of 40 seconds. Each block was preceded by a short instruction text which was shown for four seconds, otherwise a fixation cross was displayed during tasks.

Attention has previously been shown to modulate the fMRI activation on the S1 (Johansen-Berg et al. 2000; Hämäläinen et al. 2002; Nelson et al. 2004; Sterr et al. 2007; Puckett et al. 2017). Therefore, in addition to the passive task, we included a tactile oddball task in our fMRI mapping procedure. After the passive task blocks, the subjects performed a short oddball training block consisting of 15 stimuli. Next, five oddball task blocks (25 trials per block, 1200-1500 ms interstimulus interval), were presented. During the oddball task, the subjects received either standard (proximal pin) or deviant (distal pin) stimuli to the tip of their right index finger (Fig. 1c), with the deviant/standard stimulus ratio 2/23. The deviant and standard stimuli were otherwise similar, except that the deviant stimulus was produced with a more distal Braille pin with respect to the standard stimulus. The subjects were instructed to press a button of a keypad (LUMItouch, Photon Control Inc., Burnaby, Canada) with their left index finger when they perceived a deviant stimulus. The task blocks were separated by rest blocks during which the subjects were instructed to fixate on a cross. During scanning the subjects were shown visual instructions on a back-projected screen visible via an angled mirror mounted on the receiving head coil.

### 2.8. Functional MRI data preprocessing and analysis

Functional MRI data were preprocessed and analyzed using Statistical Parametric Mapping (SPM12, Wellcome Trust Centre for Neuroimaging, London, UK). The preprocessing steps included: slice-time correction, realignment, coregistration, segmentation, normalization to MNI-space using 4_th_ Degree B-Spline interpolation, and spatial smoothing with a 5 mm Gaussian kernel.

First level analyses were performed separately for the passive and oddball tasks. First, we analyzed the whole brain data of passive left and right index finger stimulation. Analyses were performed for each condition (left or right) separately and contrasting left versus right hand stimulation. In the rest of the analyses, we focused on analyzing only right-hand stimulation conditions.

After the whole-brain first-level analyses, we focused on the somatosensory representations of the right index finger. According to our *a priori* hypothesis, the fMRI activity during right index finger stimulation was suspected to be located within the left postcentral gyrus. Thus, anatomical masks covering the whole left postcentral gyrus were drawn manually for each individual and normalized into MNI-space. The resulting masks (average size 2510 voxels, SEM ± 136 voxels) were used as inclusion masks in the first level analyses. Task, instructions, and motion parameters were used as regressors. In the first-level fMRI analyses, a cluster significance threshold of *P* < 0.05 (FWE-corrected) and cluster extent threshold of 10 voxels were used, except for one subject in the passive task and for another subject in the oddball task (see the Results section).

The statistically significant fMRI clusters within the left postcentral region were back-projected to the native space of each subject using the nearest neighbor interpolation. The distance calculations between nTMS and fMRI locations were performed with an Euclidean distance formula. Paired *t*-tests were used to compare the differences between the coordinates (Diekhoff et al. 2011). Individual space was used for within-subjects distance calculations and MNI-space for between-subjects calculations.

In addition, whole-brain statistical maps were obtained for each individual and these maps were used in the second level group analysis. In the second level fMRI analysis, a cluster significance threshold of *P* < 0.0001 (uncorrected) and voxel extent threshold of 10 voxels were used.

Second-level results were visualized onto ‘mni156_2009_256’ -template using MriCroGL software (Fig. 2; http://www.mccauslandcenter.sc.edu/mricrogl/home). Visualization of the first-level results was performed using Surf Ice software (Fig. 3 and 4; https://www.nitrc.org/projects/surfice/).

**Figure 2.**
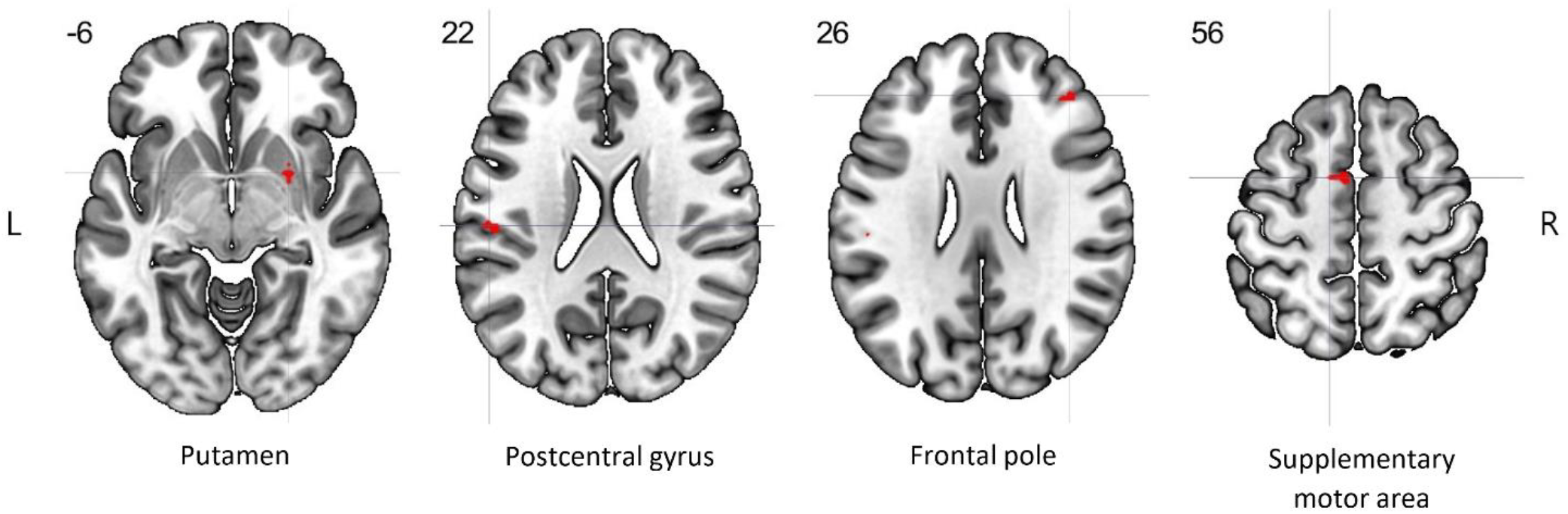
Four significant activation clusters were found in the second level fMRI analysis of the oddball task (cluster significance threshold *P* < 0.0001 [uncorrected], voxel extent threshold 10 voxels). The numbers in the upper left corner of each slice represent the z-coordinate in MNI space. The crosshairs mark the peak activations of each cluster.

**Figure 3.**
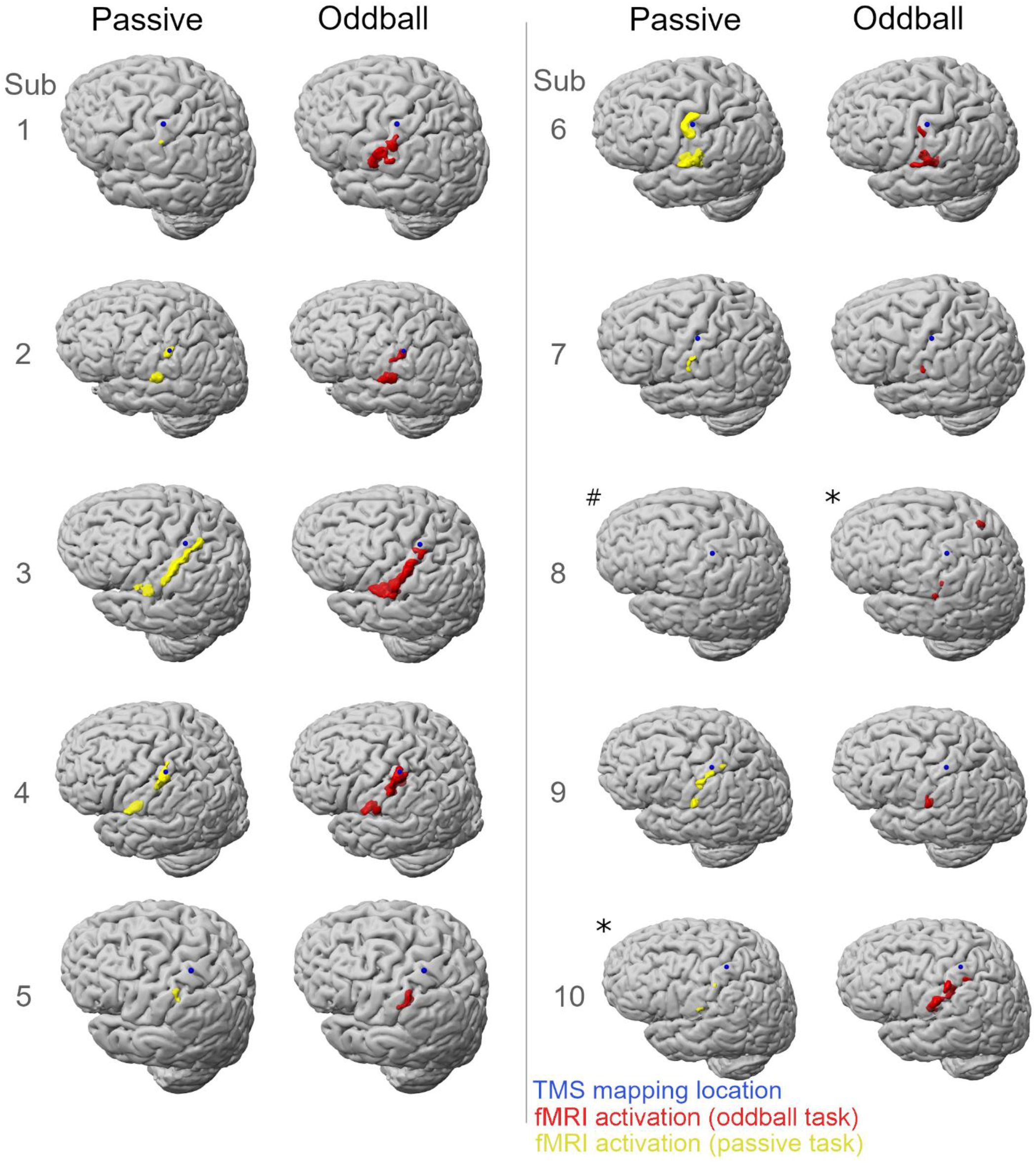
Comparison of nTMS and fMRI mapping results overlaid on individual brains. Each nTMS mapping location (S1_HS_) is marked by a blue sphere. The numbers refer to the subject numbers. Statistically significant fMRI activation clusters of the passive task are shown in yellow, and those of the oddball task in red (cluster significance threshold *P* < 0.05 FWE-corrected, cluster extent threshold 10 voxels). In two subjects, the threshold was changed (marked with asterisk; *P* < 0.001 [uncorrected], cluster extent threshold 0 voxels). Subject 8 had no significant fMRI activation in the passive task (marked with #).

**Figure 4.**
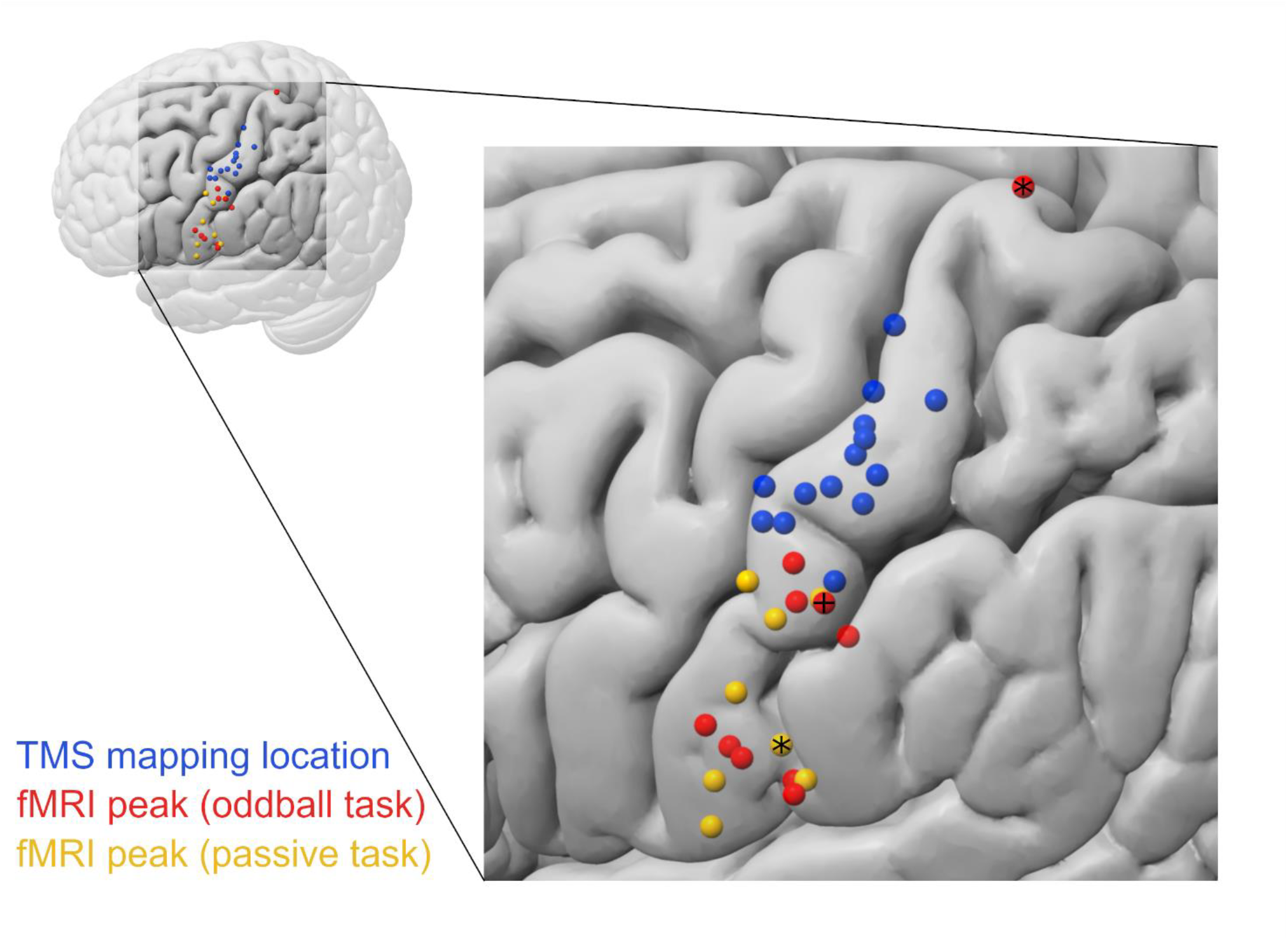
Interindividual variability of the somatosensory representations, overlaid on template brain. Blue spheres = nTMS mapping locations; red spheres = fMRI peak voxel locations of the oddball task; yellow spheres = fMRI peak voxel locations of the passive task. In one subject, the peak voxel location of the passive task was the same as the peak voxel of the oddball task (subject 1, marked with +). In both fMRI tasks, the cluster significance threshold and cluster extent threshold had to be lowered for one subject (marked with asterisks, subjects 8 and 10 in Fig. 3).

### 2.9. Statistical analyses of nTMS data

The responses (hits, misses, false alarms and correct rejections) of the nTMS experiments were analyzed with the signal detection theory -based Relative Operating Characteristics (ROC) analysis (Swets 1973), which considers the underlying psychological factors affecting subjects’ answers (see Supplementary Information). In the nTMS experiment, the area under the ROC curve (ROC[AUC]) was used as an index of response accuracy. The normality of nTMS behavioral data (ROC[AUC] -values, response times and confidence ratings) was explored with Shapiro-Wilk normality test. ROC(AUC) data had skewed distributions (skewness > -1) and did not pass the Shapiro-Wilk normality test. Thus, ROC(AUC) results were analyzed with Kruskall-Wallis tests followed by Dunn’s *post hoc* test. Response time and confidence rating data were suitable for parametric testing, so ordinary one-way analyses of variance (ANOVAs) were used for those. Behavioral data of the spatial discrimination experiment were analyzed with paired t-tests. A *P*-value of < 0.05 was considered to represent a statistically significant difference.

## 3. Results

### 3.1. Behavioral results

In the nTMS blocking experiment we investigated how navigated, single-pulse nTMS applied to the S1 affects tactile perception. For three subjects, the S1_HS_ was not found, and these subjects were not further studied. For subjects for which S1_HS_ was found (N = 14), Kruskall-Wallis test showed a significant difference between ROC(AUC) –values of the S1_HS_, the S1_NHS_ and control site conditions (*P* < 0.0001). *Post hoc* analysis showed that nTMS of the S1_HS_ impaired the tactile perception ability when compared to stimulation of control site (Oz electrode position; *P* < 0.001) or S1_NHS_ (*P* < 0.001) stimulation, while the difference between the S1_NHS_ and the cortical control site stimulation was non-significant (*P* > 0.99). ANOVA of response times showed no significant difference between the S1_HS_, the S1_NHS_ and control site conditions (*F*_2,36_ = 0.90, *P* = 0.41). Moreover, ANOVA of the confidence ratings was non-significant (*F*_2,36_ = 2.40, *P* = 0.10).

The mean ROC (AUC) in the spatial discrimination task was 0.86 (SEM ± 0.041) during nTMS of S1_HS_, and 0.87 (SEM ± 0.027) during control site (Vertex) stimulation. The difference between S1_HS_ and vertex stimulation in the spatial discrimination task was non-significant (*t*_10_ = 0.20, *P* = 0.85), suggesting that S1_HS_ was not critical for tactile spatial discrimination.

The average response accuracy in the oddball task of the fMRI experiment was 99.6 %.

### 3.2. TMS mapping results

The nTMS blocking locations varied between individuals, the largest distance between the S1_HS_ sites being 36 mm (Fig. 4). Within subjects, the average distance between the S1_HS_ and the M1_HS_ was 12 mm (SEM ± 1 mm) and the distance between the S1_HS_ and the S1_NHS_ was 11 mm (SEM ± 1 mm).

### 3.3. fMRI mapping results

In the fMRI mapping experiment, a passive task and an oddball task were used. Contrasting passive left versus right hand blocks did not show any significant activations for four subjects (for details, see Supplementary Table 1). Thus, we focused on the fMRI conditions of the right-hand stimulation. In both right-hand fMRI conditions (passive and oddball), the size, shape and location of the fMRI activation clusters varied considerably between subjects. In subject 10, in the passive task, and in subject 8, in the oddball task, there were no significant fMRI clusters when the FWE-corrected cluster significance threshold was used. Therefore, in those conditions, the cluster significance threshold was lowered to *P* < 0.001 (uncorrected) and cluster extent threshold to 0 voxels.

In the passive tactile task, the overall sizes of significant clusters varied between subjects from 6 to 389 voxels (average 108 voxels, SEM ± 41 voxels). Between subjects, the largest distance between the peak voxel locations of the passive task was 39 mm. One subject did not have any significant activation in the passive task (subject 8 in Fig. 3).

In the oddball task, the amount of significantly activated voxels varied between subjects from 14 to 450 voxels (average 148 voxels, SEM ± 44 voxels). The locations of the maximal fMRI activation (peak voxel) also varied between individuals, the largest distance between the peaks being 84 mm.

The distance between the peak voxel in the passive task and the peak voxel in the oddball task varied from 0 mm to 34 mm (calculated in native space; average 15 mm, SEM ± 4 mm). The peak voxel MNI-coordinates between the two fMRI tasks did not differ significantly in either anterior-posterior (A-P) direction (paired *t*-test: *t*_8_ = 0.80, *P* = 0.45), medial-lateral (M-L) direction (*t*_8_ = 0.52, *P* = 0.62) or inferior-superior (I-S) direction (*t*_8_ = 0.19, *P* = 0.85). Within all nine subjects who had significant activation in both tasks, the significantly activated clusters overlapped between the two tasks (average amount of overlap 65 voxels, SEM ± 23 voxels; Supplementary Table 2). There was no statistically significant difference in the cluster sizes between the two tasks (paired *t*-test: *t*_8_ = 0.91, *P* = 0.39).

In the second-level analysis of the passive and oddball tasks, there were no significant activation clusters when FWE-corrected cluster significance threshold was used. Therefore, the threshold was changed from FWE-corrected to uncorrected (*P* < 0.0001). In the second level analysis of the passive task, no significant clusters were found when using the cluster significance threshold of *P* < 0.0001 (uncorrected) and cluster extent threshold of 10 voxels. In the oddball task, four significant activation clusters were found (Fig. 2 & Supplementary Table 3).

### 3.4. Comparison of nTMS and fMRI mapping locations

Next, we investigated the correspondence between the somatosensory fMRI activations and the nTMS mapping locations. Taken together, the individual fMRI results showed activations within the left postcentral region corresponding to S1/S2 areas.

In the passive tactile task, the within-subject distances between the S1_HS_ and the nearest significantly activated fMRI voxel varied from 1 to 22 mm (average 10 mm, SEM ± 2 mm). Within subjects, the distance between the fMRI peak voxel of the passive tactile task and the S1_HS_ varied from 7 mm to 51 mm (average 26 mm, SEM ± 5 mm). The distance of the average S1_HS_ location (x = -40, y = -23, z = 56) and average fMRI peak voxel location of the passive task (x = -57, y = -17, z = 31) was 36 mm (Fig. 4 and Table 1, computed in MNI-space). The coordinates of the S1_HS_s and the peak voxels of the passive task were not significantly different in A-P direction (paired *t*-test: *t*_8_ = 1.95, *P* = 0.09). In M-L direction (*t*_8_ = 7.09, *P* = 0.0001) and in I-S direction (*t*_8_ = 3.80, *P* = 0.005), the coordinates of the S1_HS_s and peak voxels of the passive task differed significantly. The fMRI peak voxels of the passive task appeared more infero-laterally than the S1_HS_s.

**Table 1.**
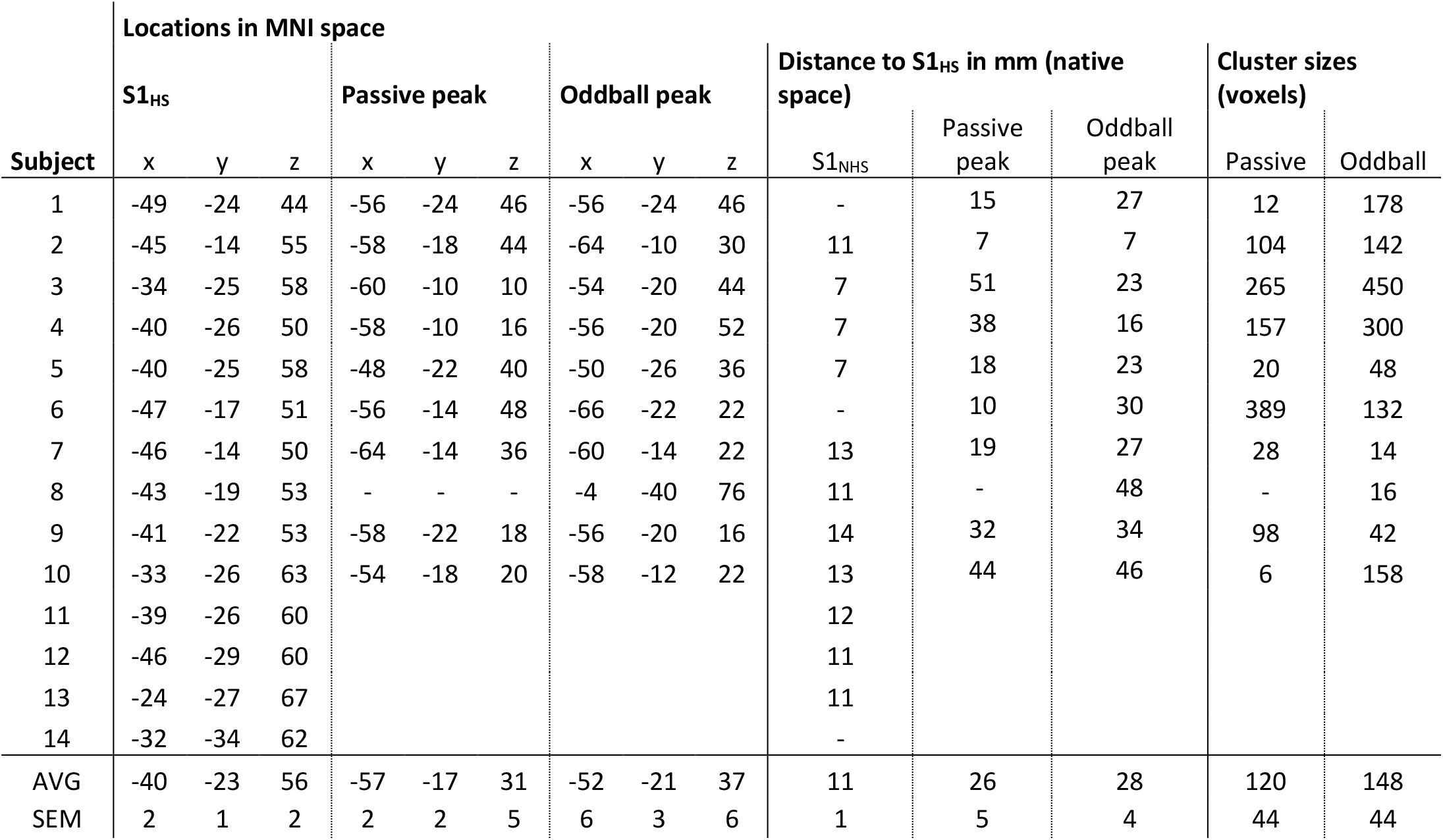
Summary of the results.

| Subject | Locations in MNI space |  |  |  |  |  |  |  |  | Distance to S1<sub>HS</sub> in mm (native space) |  |  | Cluster sizes (voxels) |  |
| --- | --- | --- | --- | --- | --- | --- | --- | --- | --- | --- | --- | --- | --- | --- |
|  | S1<sub>HS</sub> |  |  | Passive peak |  |  | Oddball peak |  |  | S1<sub>NHS</sub> | Passive peak | Oddball peak | Passive | Oddball |
|  | x | y | z | x | y | z | x | y | z |  |  |  |  |  |
| 1 | -49 | -24 | 44 | -56 | -24 | 46 | -56 | -24 | 46 | - | 15 | 27 | 12 | 178 |
| 2 | -45 | -14 | 55 | -58 | -18 | 44 | -64 | -10 | 30 | 11 | 7 | 7 | 104 | 142 |
| 3 | -34 | -25 | 58 | -60 | -10 | 10 | -54 | -20 | 44 | 7 | 51 | 23 | 265 | 450 |
| 4 | -40 | -26 | 50 | -58 | -10 | 16 | -56 | -20 | 52 | 7 | 38 | 16 | 157 | 300 |
| 5 | -40 | -25 | 58 | -48 | -22 | 40 | -50 | -26 | 36 | 7 | 18 | 23 | 20 | 48 |
| 6 | -47 | -17 | 51 | -56 | -14 | 48 | -66 | -22 | 22 | - | 10 | 30 | 389 | 132 |
| 7 | -46 | -14 | 50 | -64 | -14 | 36 | -60 | -14 | 22 | 13 | 19 | 27 | 28 | 14 |
| 8 | -43 | -19 | 53 | - | - | - | -4 | -40 | 76 | 11 | - | 48 | - | 16 |
| 9 | -41 | -22 | 53 | -58 | -22 | 18 | -56 | -20 | 16 | 14 | 32 | 34 | 98 | 42 |
| 10 | -33 | -26 | 63 | -54 | -18 | 20 | -58 | -12 | 22 | 13 | 44 | 46 | 6 | 158 |
| 11 | -39 | -26 | 60 |  |  |  |  |  |  | 12 |  |  |  |  |
| 12 | -46 | -29 | 60 |  |  |  |  |  |  | 11 |  |  |  |  |
| 13 | -24 | -27 | 67 |  |  |  |  |  |  | 11 |  |  |  |  |
| 14 | -32 | -34 | 62 |  |  |  |  |  |  | - |  |  |  |  |
| AVG | -40 | -23 | 56 | -57 | -17 | 31 | -52 | -21 | 37 | 11 | 26 | 28 | 120 | 148 |
| SEM | 2 | 1 | 2 | 2 | 2 | 5 | 6 | 3 | 6 | 1 | 5 | 4 | 44 | 44 |

In the oddball task, the within-subject distances between the S1_HS_ and the nearest significantly activated fMRI voxel varied from 4 to 29 mm (average 14 mm, SEM ± 3 mm). Within subjects, the distance between the fMRI peak voxel of the oddball task and the S1_HS_ varied from 7 mm to 48 mm (average 28 mm, SEM ± 4 mm). The distance of the average S1_HS_ location and average oddball peak voxel location (x = -52, y = -21, z = 37) was 39 mm (Fig. 4 and Table 1, computed in MNI-space). Between subjects, the coordinates of the S1_HS_s and oddball peak voxels were not significantly different in A-P direction (paired *t*-test: *t*_9_ = 0.14, *P* = 0.89) or in M-L direction (*t*_9_ = 1.85, *P* = 0.09). In I-S direction, the coordinates of the S1_HS_s and oddball peak voxels differed significantly (*t*_9_ = 2.65, *P* = 0.03). The fMRI activation maxima appeared more infero-laterally than the S1_HS_s.

The distance between the S1_HS_ and the passive peak voxel, versus the distance between the S1_HS_ and the oddball peak voxel, was statistically non-significant (paired *t*-test: *t*_8_ = 0.02, *P* = 0.98). The distance between the S1_HS_ and the nearest significantly activated voxel in the passive task, versus the distance between the S1_HS_ and the nearest significantly activated voxel in the oddball task, was also statistically non-significant (paired *t*-test: *t*_8_ = 0.80, *P* = 0.45; Supplementary Table 4). When comparing the S1_HS_ locations with the group level fMRI activation, the distance between the average S1_HS_ location and the fMRI activation maximum of the left postcentral cluster was 26 mm. Summary of the results (coordinates, distances between S1_HS_s and fMRI peak voxels, and cluster sizes) is presented in Table 1.

## 4. Discussion

Our study investigated somatosensory representations of the right index fingertip using both navigated TMS and fMRI approaches. The results revealed three key findings: (1) considerable inter-individual variability in S1 representation locations with both methods; (2) spatial discrepancies between nTMS-defined functional hotspots and fMRI peak activations within individuals; and (3) greater variability in fMRI peak activation locations compared to nTMS-defined hotspots across subjects.

The observed inter-individual variability in somatosensory representations aligns with previous invasive (Penfield and Boldrey 1937) and non-invasive (Gogulski et al. 2017) mapping studies, and neuroimaging findings (van Westen et al. 2004; Martuzzi et al. 2014). The degree of variability—up to 36 mm for nTMS hotspots, 39 mm for passive fMRI task peaks, and 84 mm for oddball fMRI task peaks—underscores the importance of individual-specific mapping approaches rather than relying on atlas- or scalp-based (Cohen et al. (1991); Seyal et al. (1997); Andre-Obadia et al. (1999)) coordinates, or coil placement posterior to the M1_HS_ (Harris et al. (2002); Morley et al. (2007); Meehan et al. (2008)). This substantial inter-individual variability likely reflects the complexity and unique organization of the somatosensory cortex in each individual (Willoughby et al. 2021). Within subjects, however, we found that the S1_HS_ was consistently reproducible and spatially distinct from the S1_NHS_, which was on average 11 mm away from the S1_HS_ and did not produce the same blocking effect (i.e. nTMS to S1_NHS_ did not block tactile perception). This spatial specificity of the blocking effect demonstrates the fine-grained functional organization of the somatosensory cortex that can be detected with appropriate mapping techniques.

A notable finding was the systematic infero-lateral displacement of fMRI peak activations relative to the nTMS-defined hotspots. This spatial discrepancy (average distance 26 mm for passive task and 28 mm for oddball task) likely reflects fundamental differences in the neurophysiological bases of these techniques. While nTMS blocking identifies cortical sites where stimulation causally disrupts tactile perception, fMRI BOLD signals capture broader hemodynamic responses that may include both primary processing and secondary network activation, potentially incorporating S2 activity (Hlushchuk and Hari 2006; Hlushchuk et al. 2015). This explanation is supported by the coordinates of fMRI activations frequently corresponding to regions associated with S2. The passive and oddball fMRI tasks produced overlapping but not identical activation patterns, with the oddball task engaging additional attentional networks that may contribute to the observed differences. Previous studies have demonstrated that attention modulates fMRI activation in S1 (Johansen-Berg et al. 2000; Hämäläinen et al. 2002), which could explain why the oddball task, requiring active monitoring of tactile stimuli, produced distinct activation patterns with even greater inter-individual variability than the passive task.

Our spatial discrimination task results further illuminate functional specificity within the somatosensory cortex. While nTMS of S1_HS_ impaired tactile detection, it did not significantly affect spatial discrimination abilities, suggesting a dissociation between detection and spatial processing pathways within S1. This functional segregation may explain why fMRI, which captures both processes simultaneously, shows activation patterns that differ from the more specific detection-related representation identified by nTMS. These findings are consistent with previous work demonstrating that S1_HS_ is critical for both detection and temporal discrimination of tactile stimuli (Hannula et al. 2005; Hannula et al. 2008), but apparently not for spatial discrimination as shown in the current study. The potential involvement of different somatosensory subregions in spatial versus temporal processing may contribute to the spatial discrepancies observed between techniques that capture different aspects of the somatosensory function. Interestingly, earlier studies using non-navigated TMS of the parietal cortex have reduced spatial discrimination ability (Porro et al. 2007; Harris et al. 2008). These findings are in line with the present results suggesting that cortical areas outside the S1_HS_ contribute to tactile spatial discrimination.

The discrepancies between nTMS and fMRI results have important methodological implications for somatosensory mapping. For functional mapping of somatosensory cortex, our findings suggest that nTMS might provide more precise localization of areas critical for tactile detection, while fMRI captures broader functional networks involved in somatosensory processing. Previous studies reporting failure to block tactile detection with S1 stimulation when targeting fMRI-defined locations (Tamè and Holmes 2016; Holmes and Tame 2019) likely reflect this spatial discrepancy between techniques. It is important to note that non-sensory factors such as changes in response criterion cannot explain the suppression of tactile detection by nTMS of S1_HS_ in our study, as only stimulation of S1_HS_ but not S1_NHS_ affected perception, despite both sites receiving identical TMS. Furthermore, our analysis using signal detection theory methods (Swets 1973) explicitly accounts for potential response bias effects, confirming that the observed blocking effect represents a genuine perceptual phenomenon rather than a cognitive bias.

The broader methodological implication of our study is that multimodal mapping approaches provide complementary information about somatosensory representations. Each technique offers distinct insights: nTMS provides causal evidence regarding cortical sites critical for specific perceptual functions, while fMRI reveals the distributed networks engaged during tactile processing. The systematic differences we observed between nTMS and fMRI localizations suggest that fMRI activations may reflect a broader activation area including both direct and indirect somatosensory processing networks, while nTMS offers a more focused method for pinpointing the functional hotspot critical for tactile detection. The spatial extent of fMRI activation clusters, which varied considerably between subjects (from 6 to 389 voxels in the passive task and 14 to 450 voxels in the oddball task), further supports this interpretation. Combined approaches may be particularly valuable in clinical contexts such as presurgical mapping, where accurately distinguishing critical functional sites from associated processing networks is essential for preservation of function.

In conclusion, our findings demonstrate substantial individual variability in somatosensory representations and systematic differences between mapping techniques. These results highlight the importance of using multimodal approaches for comprehensive functional brain mapping and underscore the need for individualized mapping in both research and clinical applications. Future studies combining nTMS with other neuroimaging methods, such as magnetoencephalography, may provide additional insights into the spatial and temporal dynamics of somatosensory processing and further refine our understanding of the complex functional organization of the human somatosensory system.

## Acknowledgements

The study was funded by The Academy of Finland (#332239), Aalto Brain Centre and Finnish Cultural Foundation. We thank personnel of AMI Centre (Aalto Neuroimaging, Aalto University School of Science, Espoo, Finland) for help in fMRI data acquisition.

## Supplementary Material

### Preliminary behavioral testing

In a preliminary experiment we studied how mechanical stimulus amplitude affects tactile perception to find optimal stimulus parameters for the TMS blocking experiment. Five subjects participated in the preliminary experiment (mean age 28 years, age range 22-35 years, 1 female), four of which also participated in the TMS experiment. Tactile stimuli were applied to the fingertip of the subject’s right index finger. To simulate the clicking sound produced by a TMS coil, a prerecorded TMS pulse sound was delivered via headphones 20 ms after the onset of the tactile stimulus.

First, a series of stimuli with a step-wise reduction in amplitude were applied to determine at what amplitude level the subject could no longer perceive the stimulus, giving a rough estimate of the tactile threshold. Subjects responded verbally whether or not they felt the tactile stimulus.

Then, because the individual tactile thresholds varied between subjects, the stimulus parameters were adjusted to obtain similar stimulus perceptions. A stimulus amplitude range, which was constant in width across all subjects, was individually shifted for each subject. The shift was performed so that the weakest stimulus in the range could not be perceived by the subject, while the strongest stimulus was clearly perceived. Ten blocks of stimuli were applied, each block consisting of 7 trials with varying tactile stimulus amplitudes within the shifted amplitude range, as well as one trial with a sham stimulus. The trials were presented in a random order. In the sham condition the prerecorded TMS pulse sound was delivered without a tactile stimulus. The rise times, and thus amplitudes, of the 7 tactile stimuli were spaced apart by 0.3 ms (approximately 10 μm rise height if the rise velocity is assumed constant). We defined the weakest stimulus in each individual subject’s amplitude range as being 1.0 in arbitrary amplitude units, and the strongest as being 2.8 arbitrary amplitude units. Subjects responded by pressing one of the two buttons of a computer mouse with their left hand. If they felt the stimulus, they responded by pressing the left mouse button, if not, they responded by pressing the right mouse button. There was no enforced response time limit, but subjects were instructed to answer as quickly and as accurately as possible. The inter-trial interval, measured from the response, was 3-3.5 seconds (randomized, step size 0.1 s).

We analyzed the data of the preliminary experiment using one-way repeated measures analysis of variance (ANOVA) with Geisser-Greenhouse correction followed by linear trend test. The results of the preliminary experiment are shown in Supplementary Fig. 1.

**Supplementary Figure 1.**
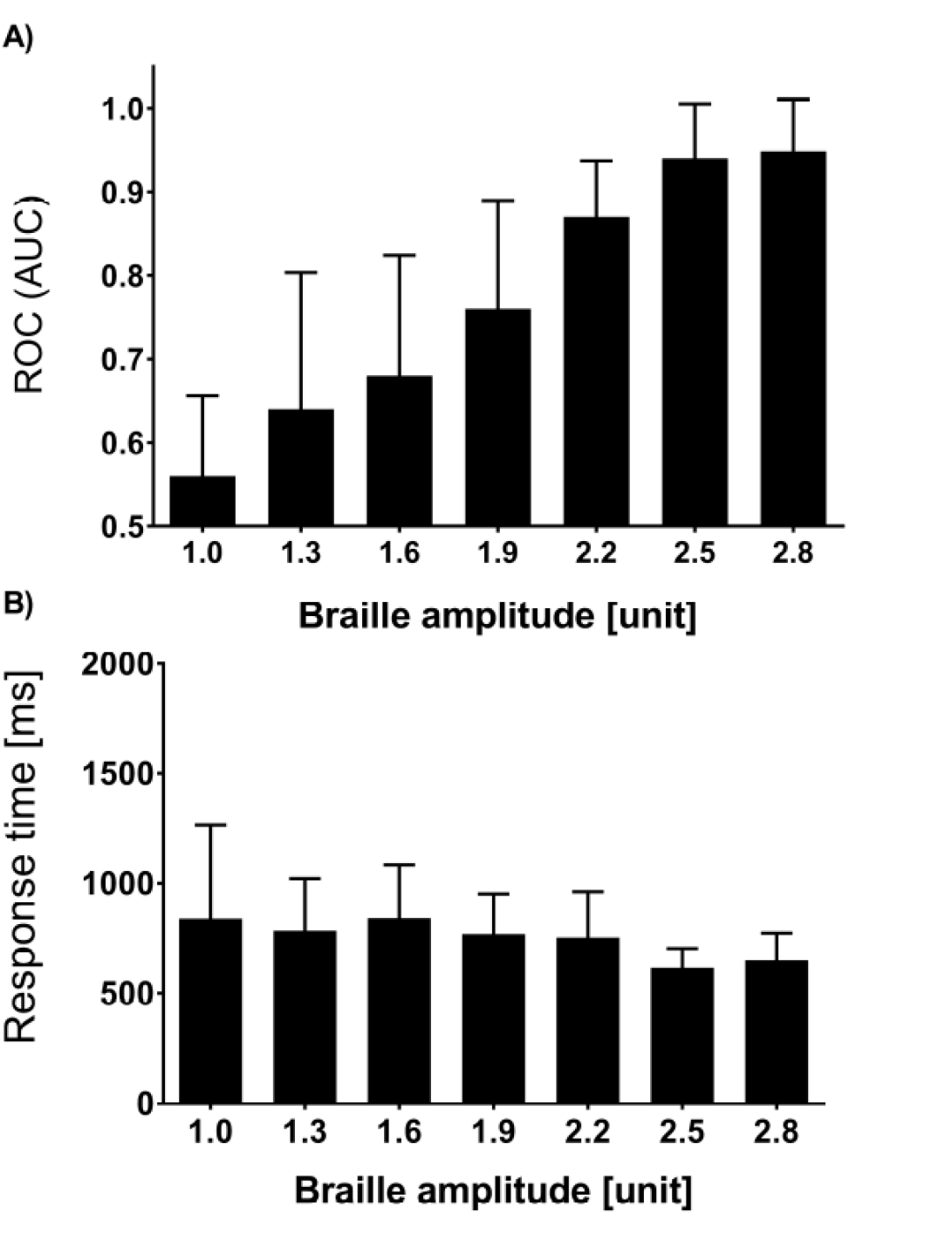
Effect of tactile stimulus amplitude on cutaneous perception. **a** The rise time of amplitude 1.0 varied between subjects in the range of 4-8.5 ms (corresponding to approximately 120-250 μm rise height if the rise velocity is assumed constant). Subjects’ ability to feel the stimuli varied significantly with stimulus amplitude (*F*_2.094, 8.376_ = 14.07, *P* = 0.002). The larger the stimulus amplitude was, the better the subjects could perceive it. Further examination showed that there was a significant linear trend (R_2_ = 0.656, *P* < 0.0001). **b** Reaction times did not vary significantly between the different stimulus amplitudes (*F*_1.519, 6.076_ = 1.385, *P* = 0.307). *Error bars* represent SD (n = 5).

**Supplementary Table 1.** fMRI first level results (contrasting passive left hand stimulation < passive right hand stimulation). The table shows only the clusters with the peak Z-value.

| Subject | Correction | k threshold | Cluster size | Peak Z-value | Peak coordinates (MNI) |  |  |
| --- | --- | --- | --- | --- | --- | --- | --- |
|  |  |  |  |  | x | y | z |
| 1 | FWE | 10 | 29 | 4.7 | -40 | -32 | -58 |
| 2 | uncor | 0 | 1 | 3.28 | -42 | -24 | 64 |
| 3 | FWE | 10 | 40 | 5.78 | -60 | -10 | 10 |
| 4 | FWE | 0 | 2 | 4.33 | -46 | -26 | 66 |
| 5 | uncor | 0 | - | - | - | - | - |
| 6 | FWE | 10 | 23 | 5.34 | -56 | -14 | 46 |
| 7 | uncor | 0 | - | - | - | - | - |
| 8 | uncor | 0 | - | - | - | - | - |
| 9 | FWE | 10 | 19 | 5.79 | -50 | -20 | 14 |
| 10 | uncor | 0 | - | - | - | - | - |

**Supplementary Table 2.** Overlapping fMRI activation between the two tactile tasks (oddball versus passive) within the region of interest.

| Subject | Overlapping voxels (MNI-space) |
| --- | --- |
| 1 | 6 |
| 2 | 88 |
| 3 | 201 |
| 4 | 135 |
| 5 | 17 |
| 6 | 87 |
| 7 | 8 |
| 8 | - |
| 9 | 38 |
| 10 | 6 |
| AVG | 65 |
| SEM | 23 |

**Supplementary Table 3.** Results of the second level analysis of the oddball task. MNI-coordinates refer to the locations of the peak voxels of each cluster. Cluster significance threshold of *P* < 0.0001 (uncorrected) and voxel extent threshold 10 voxels were used.

| Cluster region | Peak Z-value | MNI-coordinates |  |  | Cluster size |
| --- | --- | --- | --- | --- | --- |
|  |  | x | y | z |  |
| Right Putamen | 4,51 | 26 | 4 | -6 | 13 |
| Left Supplementary Motor Cortex | 4,51 | -12 | -2 | 56 | 27 |
| Left Postcentral Gyrus | 4,31 | -54 | -20 | 22 | 16 |
| Left Central Opercular Cortex |  |  |  |  |  |
| Left Supramarginal Gyrus |  |  |  |  |  |
| Right Frontal Pole | 4,27 | 40 | 40 | 26 | 16 |

**Supplementary Table 4.** Distances between the nearest significantly activated voxel and the S1_HS_, calculated in native space. The values represent millimeters.

| Subject | Distance between S1<sub>HS</sub> and nearest significant voxel |  |
| --- | --- | --- |
|  | Oddball | Passive |
| 1 | 11 | 13 |
| 2 | 5 | 1 |
| 3 | 6 | 8 |
| 4 | 4 | 4 |
| 5 | 17 | 16 |
| 6 | 7 | 4 |
| 7 | 24 | 16 |
| 8 | 24 | - |
| 9 | 29 | 7 |
| 10 | 10 | 22 |
| AVG | 14 | 10 |
| SEM | 3 | 2 |

